# Clear effects of population and sex but not rearing temperature on stress tolerance in a temperate butterfly

**DOI:** 10.1101/2024.01.12.575354

**Authors:** Nadja Verspagen, Michelle F. DiLeo, Marjo Saastamoinen

## Abstract

As temperatures are rising globally, the survival of organisms depends on their tolerance of such rising temperatures as well as resistance to indirect effects such as resource shortage under these new conditions. Genetic background and phenotypic plasticity in the form of acclimation are known to affect stress resistance, but much about genetic variation in plasticity is still unknown, especially in insects other than *Drosophila*. Here we aim to study the effect of population of origin, developmental temperature, and their interaction on stress tolerance (heat tolerance and starvation resistance). We test the beneficial acclimation hypothesis and how it is influenced by intraspecific differences. For this, we reared Glanville fritillary butterfly larvae originating from Finland and Spain at high and control temperatures, and measured their heat tolerance and starvation resistance. To assess potential costs of acclimation we also measured lifespan under control conditions. Neither adult heat tolerance nor starvation resistance were impacted by thermal conditions during development and thus we found no evidence for the beneficial acclimation hypothesis. Heat tolerance also did not differ between sex or population of origin. In contrast, we found interacting effects of population and sex on adult starvation resistance, with Spanish females outperforming other groups. Spanish females also had a longer lifespan under control conditions. Our study provides no evidence for the beneficial acclimation hypothesis but highlights the importance of population differences in stress tolerance.

**SUMMARY STATEMENT:** Despite their importance, the interacting effects of population of origin and developmental acclimation temperature on stress response have not often been studied together, especially in insects other than *Drosophila*.

## INTRODUCTION

Due to global warming the average temperature is increasing and extreme climatic events such as heatwaves are becoming more common (Easterling et al., 2000; Meehl and Tebaldi, 2004; Ragone et al., 2018; Russo et al., 2015). Insects are one of the most diverse groups of animals (Stork et al., 2015), and because they are ectotherms they are especially vulnerable to such heatwaves (Angilletta, 2009; Stange and Ayres, 2010). Their ability to survive periods of extreme heat directly depends on their heat tolerance (Bowler and Terblanche, 2008; Kingsolver et al., 2011; Klockmann et al., 2017; Radchuk et al., 2013), but also on their ability to survive other stressors occurring during heat waves such as resource shortage (Johansson et al., 2020; Maron et al., 2015). Effects of genetic background (Erić et al., 2022; Hoffmann et al., 2005; Sarup and Loeschcke, 2010) and phenotypic plasticity (Ghalambor et al., 2007; Hochachka and Somero, 2014; Leroi et al., 1994; Levins, 1969; Via et al., 1995) are known to affect stress resistance, but how plastic responses vary between individuals originating from different populations (i.e. genetic variation in plasticity) is still largely unknown. This highlights the need for studying factors affecting stress resistance in a comprehensive manner.

Intraspecific genetic differences in stress tolerance can be caused by local adaptation, resulting, for example, in environmental clines in stress tolerance (Erić et al., 2022; Hoffmann et al., 2005; Sarup and Loeschcke, 2010). Furthermore, phenotypic plasticity plays an important role in stress tolerance. The beneficial acclimation hypothesis states that heat tolerance increases after acclimation at high temperatures (Angilletta, 2009; Huey et al., 1999; Leroi et al., 1994; Somero, 2010). When acclimation happens during development, its effects can cause improved stress tolerance in the adult stages (Chidawanyika and Terblanche, 2011; Kellermann et al., 2017; Schaefer and Ryan, 2006), a form of carry-over effects (Moore and Martin, 2019). An important mechanism for this beneficial effect of developmental acclimation is the overexpression of heat shock proteins upon exposure to mild heat stress (Rinehart et al., 2006; Sejerkilde et al., 2003; Yocum et al., 1991), although this alone does not fully explain the effects of developmental acclimation on heat tolerance (Feder and Hofmann, 1999). Other contributing factors might be changes in membrane lipid composition (Hazel, 1995), cell size (Leiva et al., 2023; Verspagen et al., 2020) and metabolic rate (Hoffmann and Parsons, 1997). Besides effects on heat tolerance, developmental thermal acclimation has also been shown to increase resistance to other stressors such as starvation resistance (Pijpe et al., 2007; Scharf et al., 2015). Such cross-tolerance is possibly caused by the upregulation of general stress responses that are shared across multiple stressors (Rodgers and Gomez Isaza, 2023).

The magnitude of developmental acclimation has been shown to differ between genotypes, for example, between populations across latitude (Bozinovic et al., 2011; Ghalambor et al., 2006; Janzen, 1967). An acclimatory response might also induce fitness costs, or negative indirect carry-over effects (Krebs and Feder, 1998; Moore and Martin, 2019), which may include a reduction in activity (MacLean et al., 2017), larval viability (Loeschcke et al., 1994) or fecundity (Cao et al., 2018). Therefore it is expected that acclimation capacity increases in areas where it is beneficial, such as those at higher latitudes where thermal seasonality is high, as outlined in the latitudinal hypothesis (Bozinovic et al., 2011; Ghalambor et al., 2006; Janzen, 1967). However, both intraspecific (Brattstrom, 1970; Calosi et al., 2010; Gunderson and Stillman, 2015; Overgaard et al., 2011) and interspecific (Azevedo et al., 1998; Gilchrist and Huey, 2004; Hoffmann and Watson, 1993; Sgrò et al., 2010; van Heerwaarden et al., 2014) studies have produced results both in favour and against the latitudinal hypothesis.

In this study, we use individuals originating from two populations of Glanville fritillary butterflies from the extremes of a latitudinal cline spanning from the south of their range in Spain, where the climate is relatively warm, to the northern range edge in Finland, with a colder climate. We first aim to test whether developmental rearing temperature affects heat tolerance and starvation resistance. We hypothesize that an increase in developmental temperature will increase adult performance at both stressors, but especially under thermal stress, in line with the beneficial acclimation hypothesis (Figure 1A-D). However, scenarios of no acclimation (Figure 1E-H) or maladaptive plasticity , where increased developmental temperature decreases stress tolerance, (Figure 1I-L) are also possible. We furthermore aim to test if populations have different heat tolerance and starvation resistance, and whether there are genetic differences in acclimation response (GxE) between the populations. We hypothesize that individuals from the Spanish population, where the climate is generally warmer and dryer than in Finland, are more heat tolerant and starvation resistant (Figure 1B, F and J). According to the latitudinal hypothesis we also expect higher levels of acclimation in Finnish compared to Spanish populations as indicated by a GxE effect (Figure 1C, J and K). Population and GxE effects might occur simultaneously, resulting in both higher stress tolerance in Spain and more acclimation in Finland as hypothesized before (Figure 1D, H and L). Finally, despite potential positive effects of developmental acclimation on stress tolerance, development under mild chronic heat stress might be costly and thus lead to negative effects on other parts of life history such as larval development time, adult size and lifespan under standard conditions (Figure 1I-L). Our final aim is to test for such negative aspects, or costs, of mild chronic developmental heat stress and for this we measure developmental traits and lifespan under control conditions. Studies combining the effects of environment and population on both heat tolerance and starvation resistance across a latitudinal cline have rarely been done, especially in insects other than *Drosophila*.

**Figure 1.**
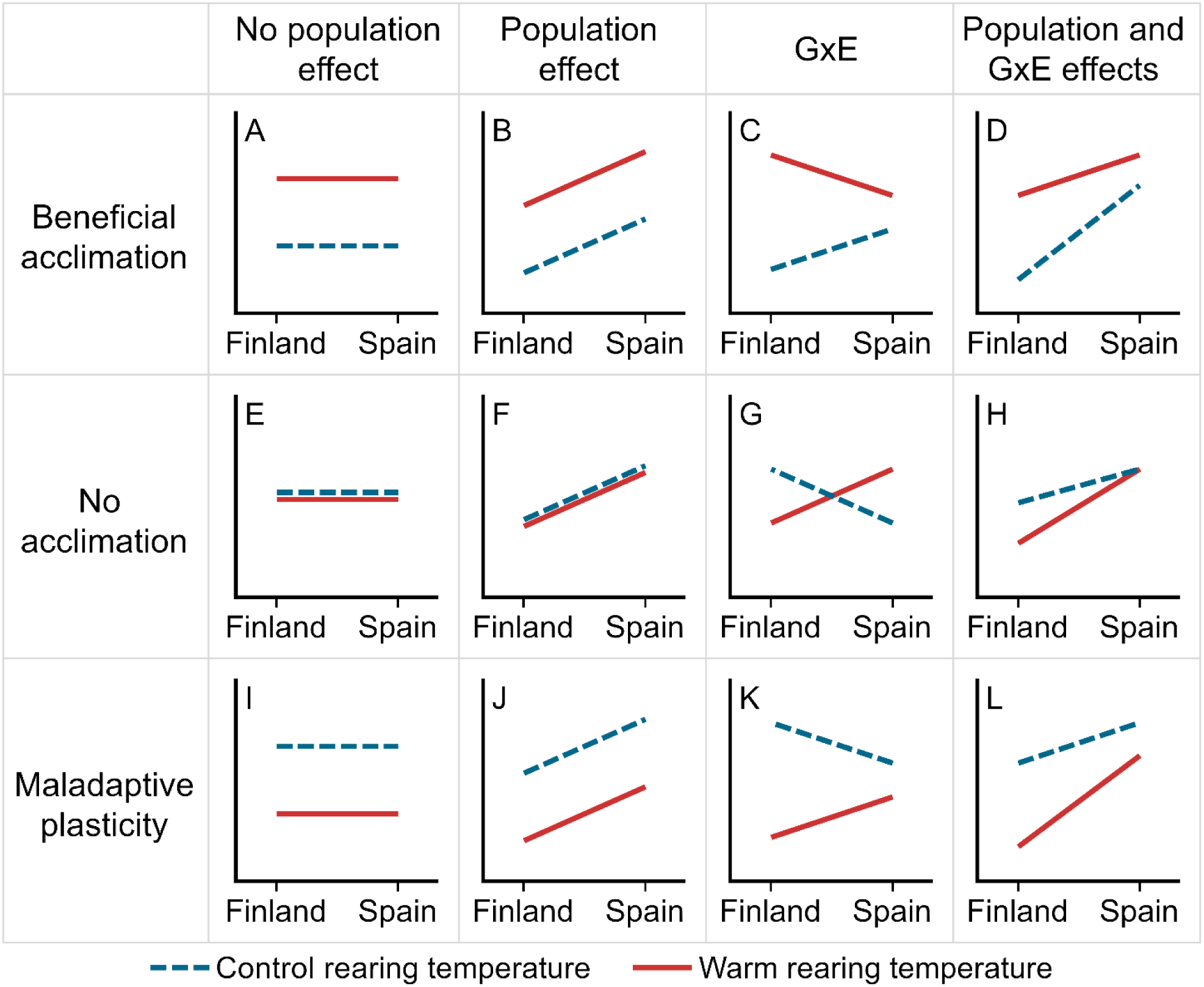
Hypothetical plots of the combined effect of developmental temperature, population and their interaction. A-D show cases of beneficial acclimation, where a warmer rearing temperature (red solid line) has a positive effect on performance compared to control rearing temperature (blue dotted line). E-H show no effect of acclimation, while I-L show situations where a higher developmental temperature has a negative effect. Panels A, E and I show no effect of population, while a main effect of population effect is present in panels B, F and J, with Spain performing better than Finland. Panels C, G and K show no main effect of population, but only the interaction effect of rearing temperature and population as indicated by a different effect of rearing temperature on performance between Finland and Spain, where Finland is more plastic than Spain. Panels D, H and L show the combined effects of population and GxE, with Spain performing better than Finland and Finland showing more plasticity than Spain.

## METHODS

### 1. Animal model & experimental populations

The Glanville fritillary (*Melitaea cinxia* Linnaeus 1758) is a temperate butterfly with a range stretching from the south of Finland until the north of Morocco, and although this may differ locally, it is not considered threatened across Europe (Maes et al., 2019). It generally adopts a univoltine lifecycle with a larval diapause, although some bivoltine populations exist in the southern range edge. Females lay egg clutches of 100-200 eggs (Saastamoinen, 2007), and pre-diapause larvae hatch and live gregariously on their host plant (Mostly *Plantago lanceolata* and *Veronica spicata*, although other *Plantago* and *Veronica* species are also used occasionally). Near the end of the summer, larvae spin a communal nest in which they overwinter until the following spring. Post-diapause larvae become solitary and can move further distances in search for food (Kuussaari et al., 2016).

In this study we used laboratory populations of the Glanville fritillary butterfly originating from Finland (Åland islands) and Spain (Catalonia). The populations have spent at least one generation in the laboratory. No ethical regulations apply for working with this species in Finland, where the experiment was conducted, and the necessary permits for collecting the individuals to start laboratory populations were present. The F0 generation from Finland was collected as diapausing larvae in September 2020 and kept under standard diapausing conditions (5°C in complete darkness with 80% relative humidity) until the following spring, when larvae were reared under standard conditions (28°C during the day and 15°C during the night, with a 12:12h light:dark cycle) and mated. Offspring was reared in family groups under standard conditions until diapause, which started in late April 2021. This F1 generation was used in the experiment. Spanish larvae were collected as egg clutches from 10 females in spring 2019, upon which the hatched larvae were reared in family groups until adulthood, and mated in spring 2020. The offspring was again reared in standard conditions and mated in spring 2021, and the resulting F2 generation was used in this experiment. Climate chambers used for rearing are Sanyo MLR-350 or Sanyo MLR-351 (Ora, Gunma, Japan), and for diapause are Rumed 4101 (Laatzen, Germany).

### 2. Larval & adult rearing

Larvae from eight families per population were taken from diapause conditions roughly 5.5 months after the initiation of diapause and kept in family groups at 25°C daytime temperature with 8°C night temperature for three days, allowing them to slowly acclimate to increased temperatures. After three days, alive larvae were weighed and moved into rearing boxes of seven larvae of the same family to avoid overcrowding. For each family, three rearing boxes were placed in standard rearing conditions (28°C during the day, 15°C during the night with a 12:12h light:dark cycle) and three were placed in hot conditions (34°C during the day and 21°C during the night, with the same 12:12h light:dark cycle), resulting in 21 larvae per family divided across three replicates per temperature treatment. Standard rearing temperature was chosen to reflect the microclimatic conditions larvae experience in the field, where temperatures can be up to 20°C above ambient air temperature (*unpublished data*, Bennett et al., 2015). Previous research has shown that larvae develop normally at 34°C with a night time temperature of 8°C (Verspagen et al., 2023) and thus here we increased the night-time temperature to impose chronic mild heat stress without crossing thermal limits. Larvae were fed daily, *ad libitum*, with leaves of the dominant host plant *Plantago lanceolata* and the rearing boxes were sprayed with water after feeding.

At pupation, pupae were weighed (Mettler Toledo XS105 DualRange, Columbus, Ohio, United States) and placed in an individually labelled cup and kept at the same temperature treatment as during the larval stage. Besides pupal mass, we measured the development time from the start of temperature treatments to pupation, and divided the natural logarithm of pupal mass by development time to calculate larval growth rate. These traits, together with adult lifespan under control conditions, were used to explore the potential costs of developmental temperature treatments. At eclosion, butterflies were sexed and marked, and assigned to either of three adult treatments: control, resource limited (to test starvation resistance) or heat tolerance by lottery draw, which ensured random but equal distribution of larvae across the treatments. All adults were kept in single-sex hanging cages divided by treatment at a constant temperature of 25°C. Butterflies from the control and heat tolerance treatments were provided with *ad libitum* 20% honey-water provided in a sponge in a petri-dish. The sponge was cleaned and refilled with honey-water every morning. Butterflies from the resource limited treatment were provided with a sponge of water that was renewed daily in the morning. On the third day after eclosion, resource limited butterflies were taken into a separate cage and provided with a sponge of 20% honey-water for six hours, after which they were returned to their original cage with water only. Every morning, dead butterflies from the control and resource limited treatments were removed and their lifespan was recorded. See figure S1A for an outline of the experimental setup with final sample sizes after mortality in larval or pupal stage.

### 3. Heat tolerance assays

Butterflies assigned to the heat tolerance treatment were tested on the morning of the third day after eclosion. At 17:00 on the day before the heat-shock assay, butterflies were taken from their rearing cage and placed in a cup with a mesh lid at room temperature in darkness, and a wet piece of cotton was placed on top of the lid until the start of the trial. The heat tolerance setup (Figure S2) consisted of a plexiglass water bath that was kept at 45°C with a heating pump, in which six 0.5 L glass cups with a see-through lid were placed. Each cup was equipped with a thermometer to measure temperature inside the cup. Temperature within the cup was kept stable at 45°C ± 1°C. At the start of the heat shock assay, a randomly selected butterfly from the day’s batch was added to one of the cups and a timer was started. When the butterfly did not move for 30 seconds, it was disturbed by quickly opening the cup and poking it with the thermometer. When this did not elicit a reaction in the form of movement, knockdown time was recorded. A maximum of two butterflies were tested at the same time, and only one observer (NV) performed all the assays. In some trials, temperature exceeded the set range, or the butterfly escaped the jar. These trials were discarded from the data, resulting in the final sample size as shown in Figure S1.

### 4. Statistical analysis

All statistical analyses are performed in R (R Core Team, 2022). Heat knockdown time, lifespan under starvation stress and lifespan under control food conditions were all tested in separate Cox Proportional-Hazards models from the *survival* package (Therneau et al., 2023) using the *coxph* function. Developmental temperature treatment, population of origin, sex and all their interactions were included in all initial models, as well as pupal mass in all and assay starting time in heat knockdown models. We performed step-wise backward model selection based on AIC by using the *step* function (Table S1). We tested for significance of model terms by using the *anova* function, and used the *emmeans* (Lenth et al., 2022) package to compute estimated marginal means and perform Tukey post-hoc tests.

To further test the effect of developmental temperature treatment, population of origin, sex, and their interactions on heat knockdown time, lifespan under resource limitation and control food conditions, we fitted linear models for each trait using the *lm* function. Again, pupal mass was included in all initial models as well as assay starting time for heat knockdown time. We used the *step* function to perform backward step-wise model selection (Table S1). We used a type III anova to test for significance of model terms by using the *anova* function from the *car* package (Fox et al., 2023) and again used the *emmeans* (Lenth et al., 2022) package to compute estimated marginal means and perform Tukey post-hoc tests. Significance letters were computed with the *cld* function from the *multcompView* package (Graves and Dorai-Raj, 2023).

## RESULTS

### 1. Stress tolerance

Firstly, we aimed to test whether developmental temperature and population affected stress tolerance. To test heat tolerance, we performed a Cox Proportional-Hazards model on time until heat knockdown, which revealed a significant three-way interaction between rearing temperature, population and sex (P = 0.039, Figure 2, Table 1). Indeed, survival probability decreased in Spanish females reared at control temperatures, but no pairwise comparisons were significant (Table S2A). We found no main effects of temperature, population or sex nor any of their two-way interactions (P > 0.05 for all terms, Figure 2, Table 1), and thus present no support for the beneficial acclimation or latitudinal hypotheses, nor the hypothesis that Spanish would be more heat-tolerant than Finnish butterflies. We additionally assessed knockdown time at 45°C, which was not significantly affected by rearing temperature, population, sex, or any of their interactions (P > 0.05 for all terms, Figure S3A, Table 2), further confirming the results of the Cox Proportional-Hazards model. Pupal mass and starting time of the heat shock assay were included in initial models but these were removed after step-wise model selection (Table S1).

**Figure 2.**
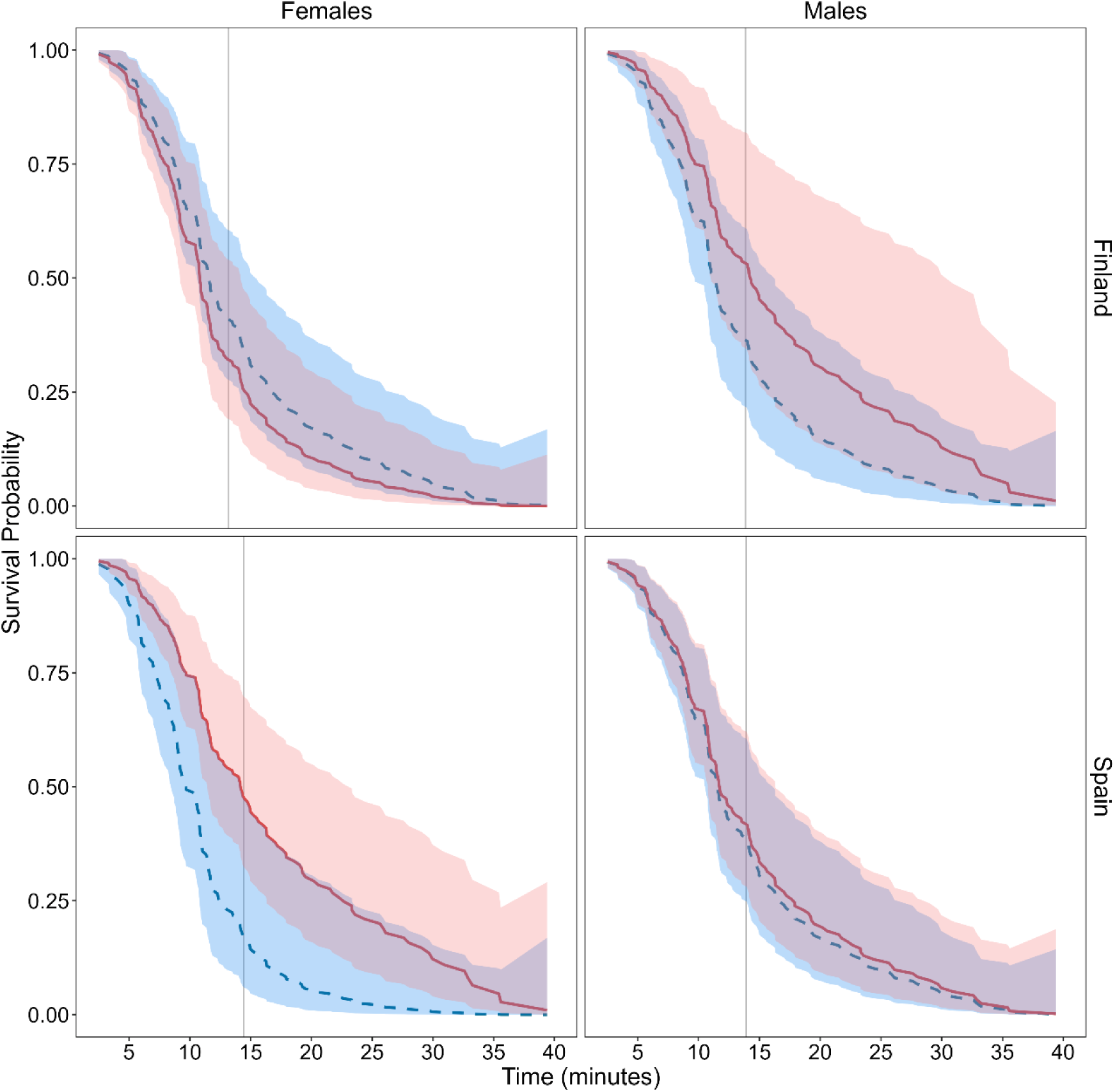
Survival probability through time at 45°C for butterflies reared under standard (blue dotted line) and hot (red solid line) conditions. Shaded area indicates the 95% confidence interval of the cox proportional hazard model. No single terms or two-way interactions were significant (P > 0.05) but we found a significant three-way interaction between temperature, population and sex. Grey vertical lines show the mean heat knockdown time for each population and sex. See Figure S1 for a breakdown of the experimental groups and sample sizes.

**Table 1.** Anova tables for Cox proportional hazard models on time until heat knockdown, time until death under food stress, and time until death under control conditions.

|  | Covariate | Loglik | Chisq | Df | P value |
| --- | --- | --- | --- | --- | --- |
| **Time until heat knockdown** | Temperature | -554.48 | 1.484 | 1 | 0.223 |
|  | Population | -554.46 | 0.029 | 1 | 0.866 |
|  | Sex | -554.32 | 0.287 | 1 | 0.592 |
|  | Temperature:Population | -553.73 | 1.174 | 1 | 0.279 |
|  | Temperature:Sex | -553.72 | 0.029 | 1 | 0.866 |
|  | Population:Sex | -553.63 | 0.174 | 1 | 0.677 |
|  | Temperature:Population:Sex | -551.49 | 4.275 | 1 | 0.039 |
| **Time until death under starvation stress** | Temperature | -649.26 | 2.306 | 1 | 0.129 |
|  | Population | -620.78 | 56.959 | 1 | < 0.001 |
|  | Sex | -611.28 | 18.990 | 1 | < 0.001 |
|  | Population:Sex | -604.55 | 13.458 | 1 | < 0.001 |
| **Time until death under control conditions** | Temperature | -670.74 | 0.008 | 1 | 0.931 |
|  | Population | -664.95 | 11.597 | 1 | 0.001 |
|  | Sex | -663.27 | 3.347 | 1 | 0.067 |
|  | Pupal.mass | -662.63 | 1.286 | 1 | 0.257 |
|  | Population:Sex | -653.54 | 18.175 | 1 | < 0.001 |

**Table 2.** Type III Anova tables for linear models run on heat knockdown time, lifespan under food stress, larval development time, pupal mass, larval growth rate and lifespan under control conditions.

|  | Covariate | Sum Sq | Df | F value | P value | R<sup>2</sup> |
| --- | --- | --- | --- | --- | --- | --- |
| **Heat knockdown time** | Temperature | 118.1 | 1 | 1.997 | 0.160 | 0.044 |
|  | Population | 3.9 | 1 | 0.066 | 0.798 |  |
|  | Sex | 6.4 | 1 | 0.107 | 0.744 |  |
|  | Temperature:Population | 93.5 | 1 | 1.580 | 0.211 |  |
|  | Temperature:Sex | 0.5 | 1 | 0.008 | 0.930 |  |
|  | Population:Sex | 9.2 | 1 | 0.155 | 0.695 |  |
|  | Temperature:Population:Sex | 159.9 | 1 | 2.702 | 0.103 |  |
| **Lifespan under starvation stress** | Temperature | 1.7 | 1 | 0.592 | 0.443 | 0.544 |
|  | Population | 291.4 | 1 | 101.787 | < 0.001 |  |
|  | Sex | 58.4 | 1 | 20.406 | < 0.001 |  |
|  | Pupal.mass | 10.0 | 1 | 3.497 | 0.063 |  |
|  | Population:Sex | 139.2 | 1 | 48.608 | < 0.001 |  |
| **Larval development time** | Temperature | 4271 | 1 | 508.312 | < 0.001 | 0.624 |
|  | Sex | 1107 | 1 | 131.781 | < 0.001 |  |
|  | Population | 1033 | 1 | 122.907 | < 0.001 |  |
|  | Temperature:Population | 60 | 1 | 7.143 | 0.008 |  |
|  | Temperature:Sex | 87 | 1 | 10.393 | 0.001 |  |
| **Pupal mass** | Temperature | 243 | 1 | 0.533 | 0.466 | 0.409 |
|  | Sex | 119546 | 1 | 262.137 | < 0.001 |  |
|  | Population | 17768 | 1 | 38.961 | < 0.001 |  |
|  | Temperature:Sex | 3473 | 1 | 7.615 | 0.006 |  |
|  | Temperature:Population | 13036 | 1 | 28.586 | < 0.001 |  |
|  | Sex:Population | 8791 | 1 | 19.278 | < 0.001 |  |
| **Larval growth rate** | Temperature | 2.724 | 1 | 621.911 | < 0.001 | 0.675 |
|  | Sex | 0.510 | 1 | 116.427 | < 0.001 |  |
|  | Population | 0.726 | 1 | 165.758 | < 0.001 |  |
|  | Temperature:Population | 0.238 | 1 | 54.326 | < 0.001 |  |
|  | Sex:Population | 0.014 | 1 | 3.230 | 0.073 |  |
| **Lifespan** | Temperature | 0.4 | 1 | 0.010 | 0.921 | 0.224 |
|  | Sex | 24 | 1 | 0.595 | 0.442 |  |
|  | Population | 608.8 | 1 | 15.099 | < 0.001 |  |
|  | Pupal.mass | 213.1 | 1 | 5.284 | 0.023 |  |
|  | Sex:Population | 1031.5 | 1 | 25.580 | < 0.001 |  |

We then tested starvation resistance by measuring lifespan under starvation stress. We again performed a Cox Proportional-Hazards model and found that Spanish larvae had higher survival probability across time with limited resources than Finnish individuals (P < 0.001, Figure 3, Tables 1 and S2B), as hypothesized. We furthermore found that females had higher survival probability than males (P < 0.001, Figure 3, Tables 1 and S2B). There was also a significant interaction between sex and population (P < 0.001, Figure 3, Tables 1 and S2B), with Spanish females having a higher survival probability than other groups. We found no effect of rearing temperature (P = 0.129, Figure 3, Tables 1 and S2B), and the interaction between temperature and population was removed during step-wise model selection (Table S1), indicating no evidence for the beneficial acclimation or latitudinal hypothesis. Linear models performed on lifespan under resource stress revealed similar trends (P < 0.001, Figure S2B, Table 2).

**Figure 3.**
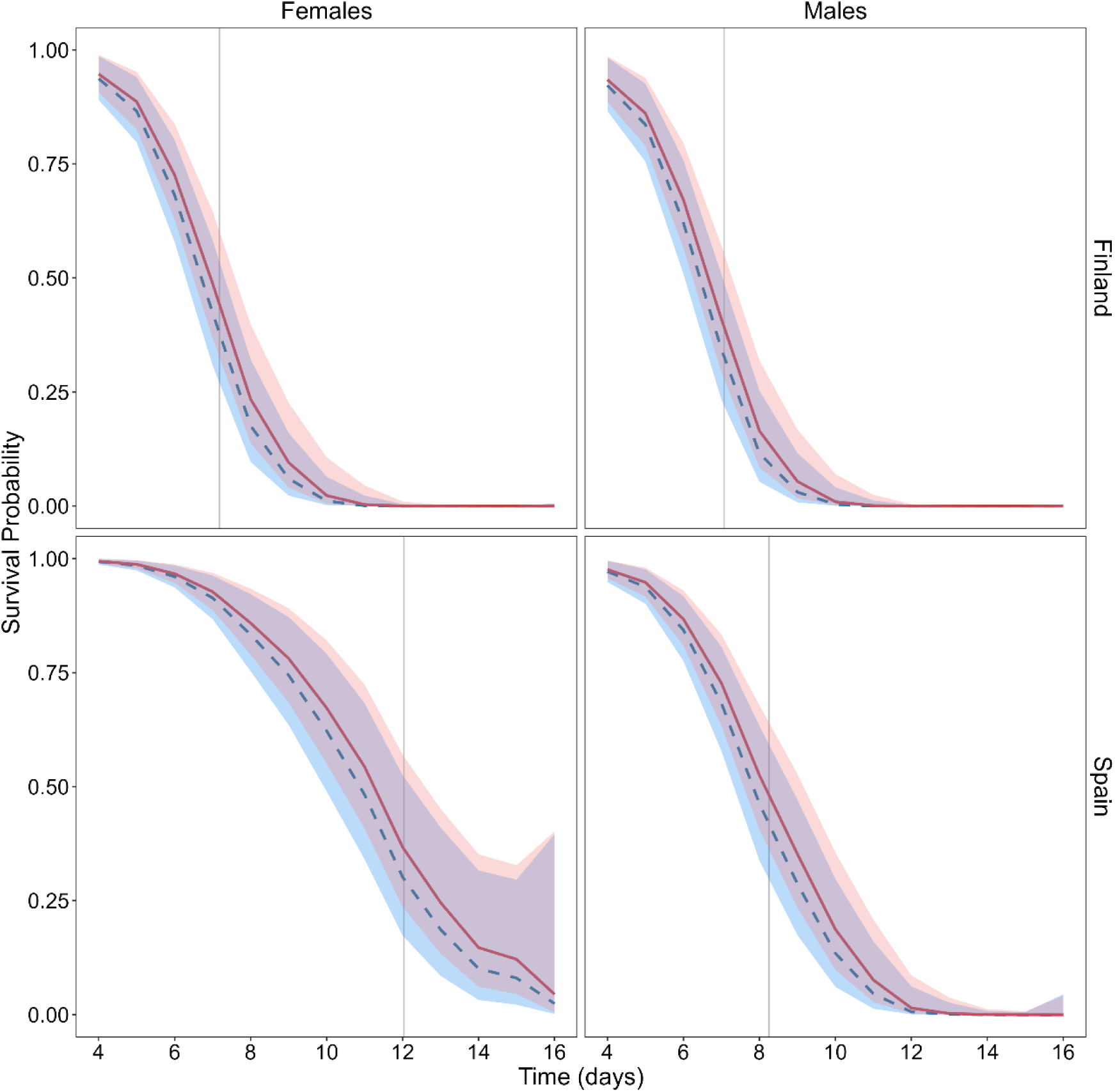
Survival probability through time under starvation stress for butterflies reared under standard (blue) and hot (red) conditions. Shaded area indicates the 95% confidence interval of the cox proportional hazard model. Spanish and female butterflies are significantly more starvation resistant (P < .001), and there is a significant interaction between population and sex (P < 0.001). Larval rearing temperature did not significantly affect starvation resistance (P = 0.129) Grey vertical lines show the mean lifespan for each population and sex. See Figure S1 for a breakdown of the experimental groups and sample sizes.

### 2. Fitness costs

To test if differences in stress tolerance of adults could be accompanied by fitness costs accrued during the larval stage, we tested whether larvae that developed under mild chronic heat stress showed differences in development time, larval growth rate, pupal mass or adult lifespan. Larvae from Spain developed significantly faster than those from Finland (P < 0.001), larvae reared in hot conditions developed faster than those reared under control conditions (P < 0.001), and males developed faster than females (P < 0.001, Figure 4A, Table 2). Spanish larvae and females in general were more responsive to temperature than Finnish larvae and males, respectively (temperature-population interaction, P = 0.008 and temperature-sex interaction, P = 0.001). Pupal mass did not differ between larvae reared at different temperatures (P = 0.466), but pupae from Spain were heavier than those from Finland (P < 0.001) and females were larger than males (P < 0.001, Figure 4B, Table 2). Furthermore, significant two-way interactions were present for all terms. While pupal mass decreased slightly with increased rearing temperature in Finnish larvae, it increased with temperature in Spanish larvae (P < 0.001). Generally females responded negatively to temperature while males respond positively (P < 0.001), and sex differences werelarger in Finland than in Spain (P < 0.001). Larval growth rate was higher in Spain than Finland (P < 0.001), increased with temperature (P < 0.001) and was higher in males than females (P < 0.001, Figure 4C, Table 2). Additionally, larvae from Spain responded more strongly to temperature than those from Finland (P < 0.001). Finally, adult lifespan under control conditions was not affected by larval rearing temperatures (P = 0.921) or sex (P = 0.442), but Spanish butterflies lived generally longer than Finnish butterflies. This effect can mainly be explained by the significant interaction between population and sex (P < 0.001), since Spanish females lived significantly longer than males but this was not the case for Finnish females. An increase in pupal mass also significantly increased lifespan (P = 0.023). We then performed Cox proportional hazard models, in which population (P = 0.001) as well as the interaction between population and sex (P < 0.001) significantly affected time until death (Figure 5, Table 1 and S2C). Similar to the analysis with linear models, Spanish females lived longer than all other groups.

**Figure 4.**
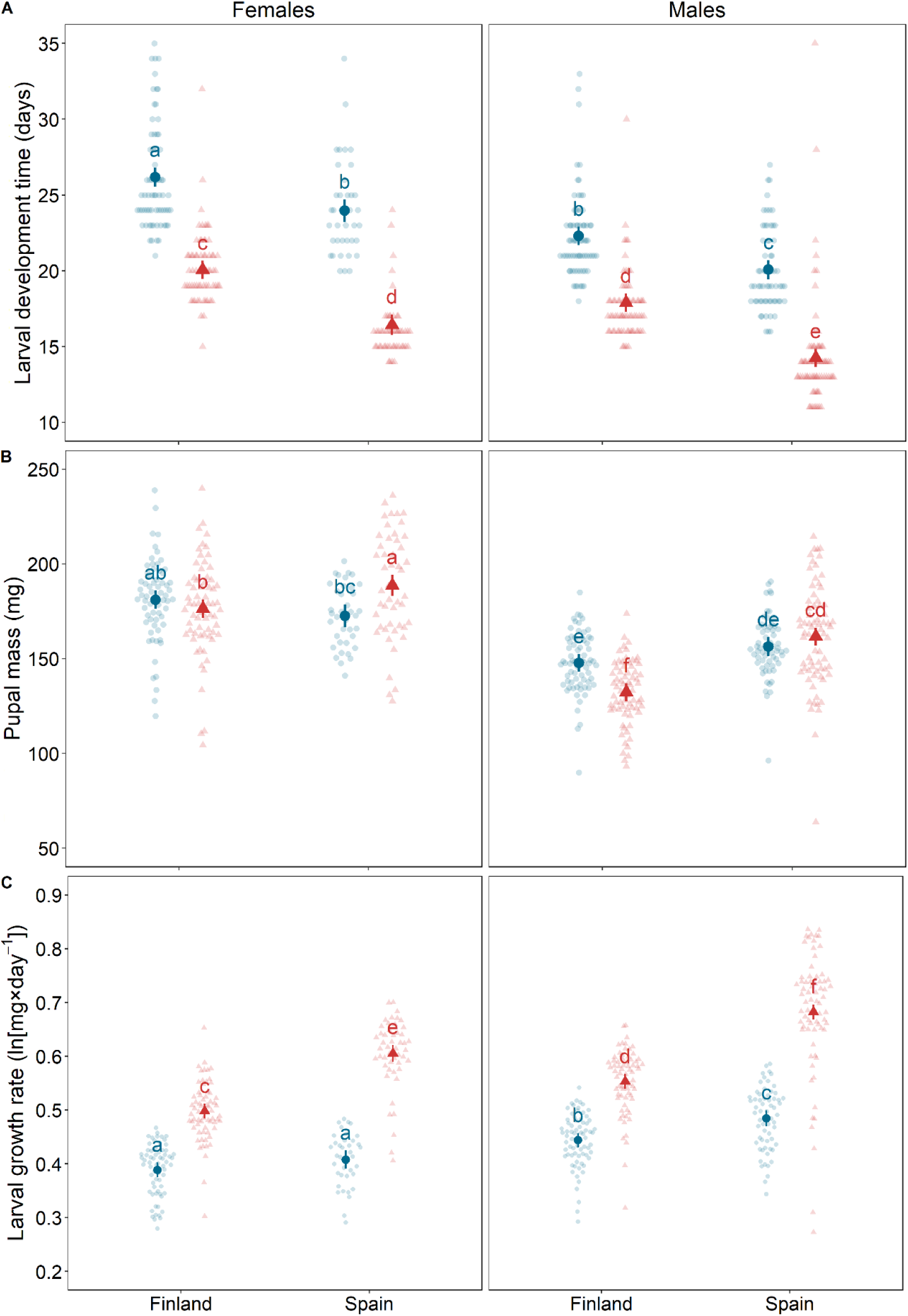
Larval development time (A), pupal mass (B) and larval growth rate (C). Blue circles indicate individuals that developed under control temperature, while red triangles show individuals that developed under hot conditions. Points indicate the estimated marginal mean±c.i. Transparent points show raw data. Letters indicate significantly different groups (P < 0.05) as determined by Tukey’s post-hoc test. See Figure S1 for a breakdown of the experimental groups and sample sizes.

**Figure 5.**
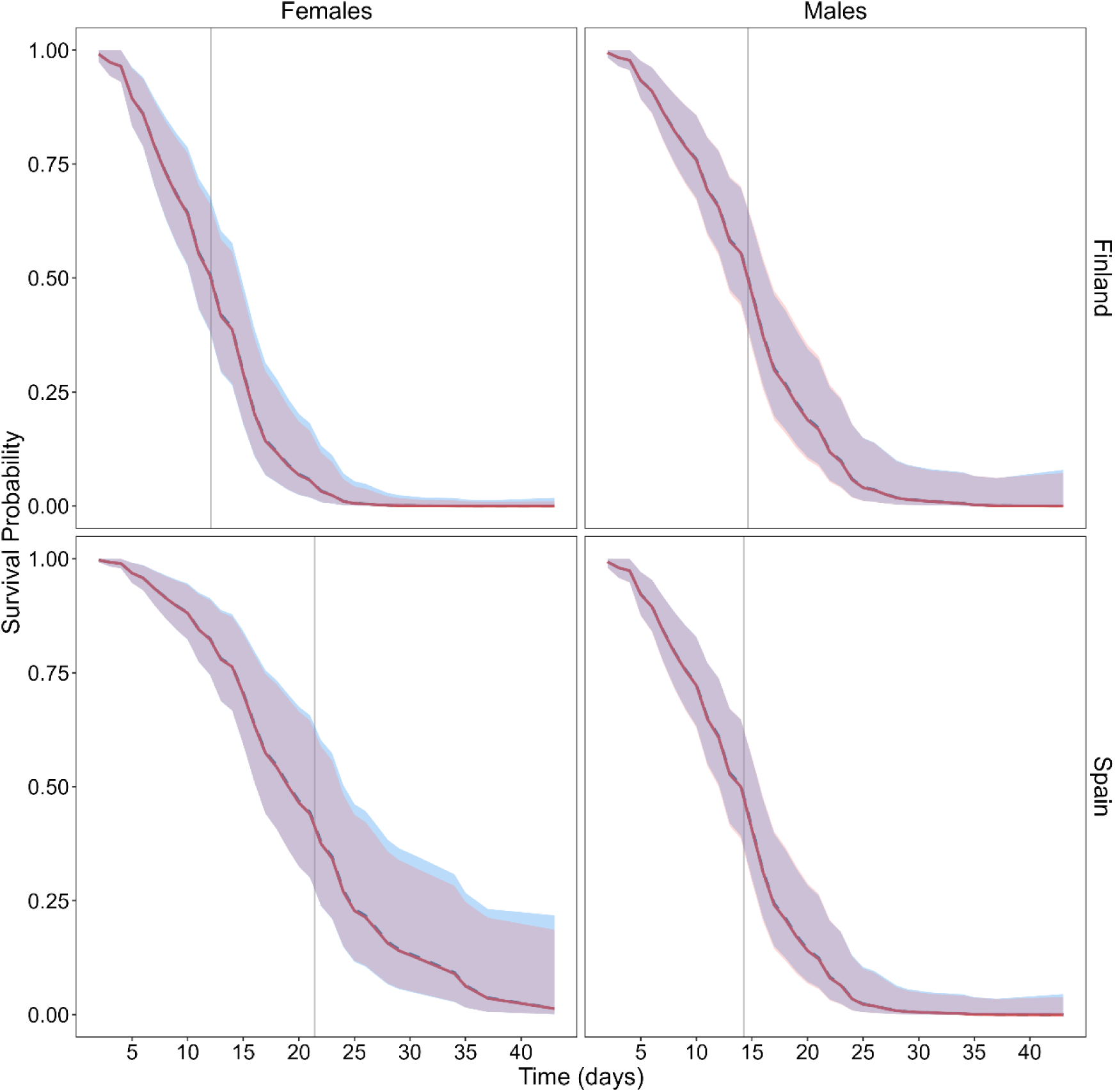
Survival probability through time under control conditions for butterflies reared under standard (blue) and hot (red) conditions. Shaded area indicates the 95% confidence interval of the cox proportional hazard model. Spanish butterflies live significantly longer than Finnish ones (P = 0.001) and we found a significant population by sex interaction (P < 0.001), showing that Spanish females live longer than other groups. Grey vertical lines show the mean lifespan for each population and sex. See Figure S1 for a breakdown of the experimental groups and sample sizes.

## DISCUSSION

One of the main aims of this study was to test for differences in stress tolerance across populations originating from different latitudes. We hypothesised that individuals from Spain would be more heat tolerant as well as more resistant to starvation due to local adaptation to originating from warmer and dryer conditions (Figure 1B, F and J or D, H and L). Spanish individuals were found to be more starvation resistant and such patterns across latitudinal clines are also present in *Drosophila* (Arthur et al., 2008; Karan et al., 1998) and other fruit flies (Weldon et al., 2018). We found that especially Spanish females were highly starvation resistant, while this effect was smaller in Spanish males or Finnish butterflies, as indicated by a significant sex by population interaction. A potential reason for this is that starvation resistance is significantly positively affected by body size (i.e. pupal mass). Spanish butterflies are larger than Finnish ones, and females are larger than males. These larger individuals could have a higher absolute resource storage, which would positively impact starvation resistance (Ballard et al., 2008; Goenaga et al., 2013). However, the difference in body size between Spanish and Finnish females is small and thus does not fully explain the observed difference in starvation resistance. Females generally live longer than males and their reproductive fitness is highly linked to lifespan, whereas this is not the case in males, who benefit from a faster pace-of life (Allen et al., 2011). We found that general lifespan was indeed also higher in Spanish females than in other groups, and lifespan has been shown to be positively correlated to starvation resistance (Lin et al., 1998; Service et al., 1985). Sexual dimorphism in starvation resistance has furthermore been demonstrated in earlier work in *Drosophila* (Ballard et al., 2008; Goenaga et al., 2010; Harbison et al., 2005; Robinson et al., 2000) and can be related to sex-specific differences in expression of reproductive traits correlated with starvation resistance such as egg production (Chippindale et al., 1993; Salmon et al., 2001) and ovariole number (Wayne et al., 2006). However, why this difference in starvation resistance with sex is not present in the Finnish population remains to be studied.

Furthermore, we hypothesized that due to living in warmer conditions, Spanish butterflies would be more heat-tolerant than Finnish ones (Figure 1B, F and J or D, H and L). Although Spanish females reared under warmer conditions do survive longer under acute heat stress than those reared at control temperature as indicated by a significant three-way interaction between population, rearing temperature and sex, no consistent pattern of beneficial acclimation was found. Even though latitudinal variation in thermal tolerance is present in certain species and populations (Hoffmann et al., 2005; Loeschcke et al., 1994; Sisodia and Singh, 2010), other studies have found no effect of latitude (Kimura, 2004; Lockwood et al., 2018). This may be related to the fact that in temperate regions, thermal variability is high and organisms thus need large thermal ranges even when average temperatures are low (Janowiecki et al., 2020). Additionally, selection on upper thermal limits might be lower in mobile species or life-stages such as adult butterflies because they are able to avoid thermal extremes through behavioural thermoregulation (Buckley et al., 2015; Dillon et al., 2009; Gunderson and Leal, 2012; Llewelyn et al., 2017; Muñoz et al., 2016). Finally, even in the presence of latitudinal selection for heat tolerance, upper thermal limits might fail to evolve due to evolutionary constrains (Hangartner and Hoffmann, 2016; Schou et al., 2014; van Heerwaarden et al., 2016). Importantly, in our study we could only include two populations and thus conclusions on latitudinal adaptation cannot be drawn with certainty. It is possible that the differences between populations we found are caused by other factors than local adaptation to heat or drought, and future studies should include more populations.

Our second research question was to study the effect of rearing temperature on adult stress tolerance, and whether there are interactions between rearing temperature and population. In line with the beneficial acclimation (Figure 1A-D) and latitudinal hypotheses (Figure 1C, G and K or D, H and L), we expected that warm-reared larvae would result in more stress tolerant butterflies and that Finnish butterflies would show more plasticity than Spanish ones. We did not find any evidence for these as there was no effect of rearing developmental temperature on either adult heat tolerance or starvation resistance, or any significant interactions between temperature and population in either stress related traits. Phenotypic plasticity is expected to be most beneficial in predictably variable environments (Chevin and Lande, 2015; Reed et al., 2010), since plastic organisms are at risk of a mismatch between environment and phenotype, leading to maladaptive plasticity, when environmental variation is not predictable (Ghalambor et al., 2007; Hoffmann and Bridle, 2022). Previous analysis from Finland, shows low predictability between spring and summer conditions (Halali and Saastamoinen, 2022). Although we do not know whether the same is true for the Spanish population, this lack of environmental predictability might explain the lack of developmental acclimation. Instead, such unpredictable environmental conditions might lead to more short-term beneficial acclimation in adult life-stages (Willot et al., 2021).

The final aim of this study was to investigate whether development under mild chronic heat stress caused negative carry-over effects (Figure 1I-L), and whether these effects would be different between populations (Figure 1). In general, increased rearing temperature did not seem costly as development time decreased and growth rate increased with rearing temperature, body size was unaffected and lifespan did not shorten with rearing temperature. This might indicate that our chosen warm rearing temperature was not high enough to elicit mild heat stress, potentially explaining the lack of acclimation response to future stress. Despite the lack of negative consequences (i.e. maladaptive plasticity or negative carry-over effects), we did find clear effects of high rearing temperature on developmental traits. Development time decreased and growth rate increased with temperature as expected, while pupal mass did not change with temperature. This is unexpected since mass is generally found to decrease with increasing rearing temperature in ectotherms according to the Temperature Size Rule (Angilletta and Dunham, 2003; Atkinson, 1994). In previous studies on the Glanville fritillary butterfly, the pre-diapause larvae from southern latitudes were shown to have canalized body size across rearing temperatures and this effect has been attributed to the lack of seasonal time constraints in these areas (Verspagen et al., 2023). Potentially, the canalization of pupal mass in both populations found in the present study indicates that time constraints are not as pressing in the post-diapause stage. Furthermore, in the current study Spanish larvae developed and grew faster than Finnish ones, especially at high temperatures, and were more responsive to temperature than Finnish larvae in these traits. This is opposite to patterns found in pre-diapause larvae of this species (Verspagen et al., 2023), where southern larvae develop slower and are less plastic in this trait than those from the north. Similarly to pupal mass, this could show that seasonal constraints are lower in post-diapause stages, but more research would be needed to find out definite causes.

To conclude, we show that stress resistance is dependent on population of origin and sex, but show no evidence for the beneficial acclimation hypothesis or for differences in acclimation response between populations as outlined by the latitudinal hypothesis. Furthermore, despite not finding evidence for costs of high rearing temperature on developmental traits or lifespan, development is highly dependent on temperature, population of origin and sex. Our study emphasizes the importance of population differences in both stress resistance as well as regular performance. Future studies testing different or more extreme developmental stressors and including more populations across a latitudinal cline will further clarify latitudinal patterns in carry-over effects of developmental stress.

## ACKNOWLEDGEMENTS

We thank Constanti Stefanescu, Alma Oksanen, Debra Dileo, Sandra Bartolini, Antoni Mariné and Josep Planes for helping to collect the butterflies that started the Spanish laboratory population, and Suvi Ikonen as well as all students involved in the Åland meadow survey for collecting the Finnish laboratory population used in this study. We also thank Morgane Frapin and Esa Pekka Tuominen for help with assembling the heat tolerance setup.

## DATA AVAILABILITY

The data and code used in this research will be made available on Dryad upon acceptance of the manuscript

